# High-Resolution 16S rRNA and Metagenomics Reveal Taxonomic, Functional Restructuring and Pathogen Persistence in Egyptian Treated Wastewater

**DOI:** 10.1101/2025.09.16.676362

**Authors:** Aya G. Atalla, Eman M. Ismail, Shaymaa Abdelmalek

## Abstract

Wastewater treatment plants (WWTPs) are pivotal for environmental protection and public health, yet the ecological dynamics of microbial communities during treatment remain poorly characterized in arid, resource-limited areas. This study presents high-resolution 16S rRNA V3–V4 amplicon sequencing and predictive metagenomics to survey bacterial populations in influent and effluent samples from five major Egyptian WWTPs. Results revealed a dramatic increase in microbial richness and evenness post-treatment, with pronounced taxonomic restructuring: copiotrophic Proteobacteria, dominant in influent streams, declined sharply, while stress-tolerant lineages including Actinobacteriota, Firmicutes, and Bacteroidota emerged in effluents. Multivariate analyses (DCA, NMDS, PCoA) confirmed consistent, significant compositional turnovers, with statistically robust separation between input and output communities. Although community diversity and functional metabolic capacity expanded, several clinically relevant genera—Escherichia, Legionella, and Campylobacter—persisted in effluents at non-trivial levels, raising concerns for water reuse safety. Functional profiling highlighted increased metabolic versatility and resilience in treated microbiomes, particularly enrichment in nutrient cycling, xenobiotic degradation, and stress adaptation pathways. Community trajectories were strongly influenced by site-specific physicochemical factors, underscoring the need for tailored surveillance and optimized operation. Comparative analyses reinforced the universality of Proteobacteria dominance in raw sewage and demonstrated widespread post-treatment diversification, yet also revealed local variability shaped by plant design and regional characteristics. Together, these findings detail the molecular ecology underpinning wastewater treatment, identify persistent hazards, and advocate for region-specific monitoring and intervention strategies aligned with sustainable water reuse and One Health imperatives.

**Importance:** This study addresses a critical knowledge gap in wastewater management within arid and developing regions by elucidating the taxonomic and functional transformations of bacterial communities during treatment. The data show that while conventional WWTPs reliably increase microbial diversity and foster the development of resilient, functionally versatile communities, conventional processes may fail to fully eliminate key pathogens, thus posing ongoing public health risks. The persistent detection of clinically relevant genera in treated effluents highlights the limitations of standard monitoring and underscores the urgent need for integrated microbiome and resistome surveillance. These insights emphasize that effective wastewater management must go beyond chemical contaminant removal and include systematic, context-sensitive biological monitoring to safeguard water reuse. The study advances microbial ecology as a cornerstone of safe, sustainable water management, providing actionable evidence for policymakers and practitioners aiming to minimize pathogen dissemination and optimize treatment outcomes.

## Introduction

Safe and sustainable reuse of wastewater is a cornerstone of modern water management in water-limited environments, underpinning solutions for water scarcity, environmental protection, and resource recovery (1–6). The World Health Organization’s (WHO) Guidelines for the Safe Use of Wastewater, Excreta and Greywater position treated wastewater as a strategic resource, supporting long-term water and nutrient security while reducing adverse environmental effects from untreated effluent (7, 8). These principles align closely with United Nations Sustainable Development Goal 6 (SDG 6), emphasizing the global imperative for effective treatment and equitable sanitation access (9, 10). Despite rapid advances in engineered treatment systems, a critical challenge persists in ensuring the effective removal of pathogenic microorganisms and harmful chemical contaminants so that the benefits of reuse are balanced with risks to human and ecosystem health (11–19).

Wastewater Treatment Plants (WWTPs) function as dynamic ecological filters, applying physical, chemical, and biological processes to transformed influent streams (3, 20–22). Each treatment stage can drive shifts in microbial community structure, sometimes reducing diversity, sometimes selecting for functional specialists with the potential to pose public health risks (23, 24). Monitoring and understanding these microbiome dynamics, especially as they relate to the emergence, persistence, or suppression of key taxa, is essential for enhancing treatment efficiency, ensuring the reliability of effluent quality, and anticipating downstream impacts (25). Gaining detailed insights into the trajectories of microbial communities through WWTPs in diverse environmental and operational contexts remains a crucial yet underexplored area of research (11).

Bacterial communities in wastewater are taxonomically and functionally diverse, encompassing lineages with roles in nutrient cycling, organic matter degradation, and pollutant transformation, as well as those with pathogenic or opportunistic potentials(26–28). These communities originate predominantly from anthropogenic sources, and their fate during treatment may have far-reaching impacts on environmental and public health upon release or reuse(29). Without robust monitoring protocols, both beneficial and potentially harmful aspects of microbial community shifts may evade detection, undermining advances in water purification technology and the sustainable management of water resources(30, 31).

In Egypt, most domestic sewage is processed via WWTPs, and recent figures indicate that up to 87.8% of collected sewage undergoes treatment(32). While biological treatment is the prevailing strategy nationwide, there are no legislative mandates for systematic microbiome monitoring within WWTPs (33). Most research in the Egyptian context has centred on chemical pollutant removal, leaving a critical gap in our understanding of microbial community composition and dynamics before and after treatment (34–38). Addressing this knowledge gap is key to optimizing microbial contributions to water quality and mitigating environmental and public health risks associated with effluent reuse or discharge.

Here, building upon prior work evaluating chemical contaminant removal (Atalla *et. al.*, under review), we provide the first comprehensive analysis of bacterial community structure in five representative Egyptian WWTPs. By combining high-throughput sequencing and robust ecological analysis, this study offers novel insights into microbial diversity shifts associated with wastewater treatment and sets the stage for advancing sustainable, safe effluent reuse strategies in arid and developing regions.

## Results

### Physicochemical results

Physicochemical analyses showed that the municipal wastewater treatment reduced organic load and nutrients across all WWTPs, with variable efficiency. Chemical Oxygen Demand (COD) dropped below 50 mg/L and Biochemical Oxygen Demand (BOD) below 30 mg/L in most effluents, except slightly higher BOD in plant 5. Total Organic Carbon (TOC) declined markedly. Total Nitrogen (TN) removal exceeded 50% generally, except for 40% in plant 4. Total Phosphorus (TP) mostly decreased post-treatment, but plant 3 showed an unexpected TP increase (Atalla *et. al.*, under review) (Table A1).

### Wastewater Treatment Drives Pronounced Microbial Community Shifts

High-throughput 16S rRNA V3–V4 amplicon sequencing generated high-quality, deeply sequenced datasets for paired influent and effluent samples from five major Egyptian WWTPs. The datasets yielded 2,154 amplicon sequence variants (ASVs) across 18 phyla, 38 families, and 52 genera, with robust sequencing coverage (Q20/Q30 >90%, mean read length ≈425 nt).

### Enhanced Richness and Diversity Following Treatment

Alpha diversity metrics revealed a striking expansion of microbial richness and evenness in effluent relative to influent communities. Rarefaction analysis demonstrated that several effluent samples (e.g., O2, O4) exceeded 6,000 observed ASVs, while all influents plateaued below 2,000; median Chao1 richness increased nearly fourfold after treatment (Input ≈ 1,693 vs. Output ≈ 6,440; Wilcoxon p = 0.045) (Fig.1) (Fig. A2). The Shannon and Simpson indices also increased (Shannon: 5.94 → 8.70; Simpson: 0.959 → 0.988), indicating not just increased species counts but more evenly distributed communities. Pielou’s evenness rose from 0.557 in influent to 0.698 in effluent, and the dominance index fell from 0.041 to 0.012. These trends demonstrate a marked reduction in highly dominant taxa post-treatment, with a broader, more balanced taxonomic profile emerging in treated waters.

**Fig. 1.**
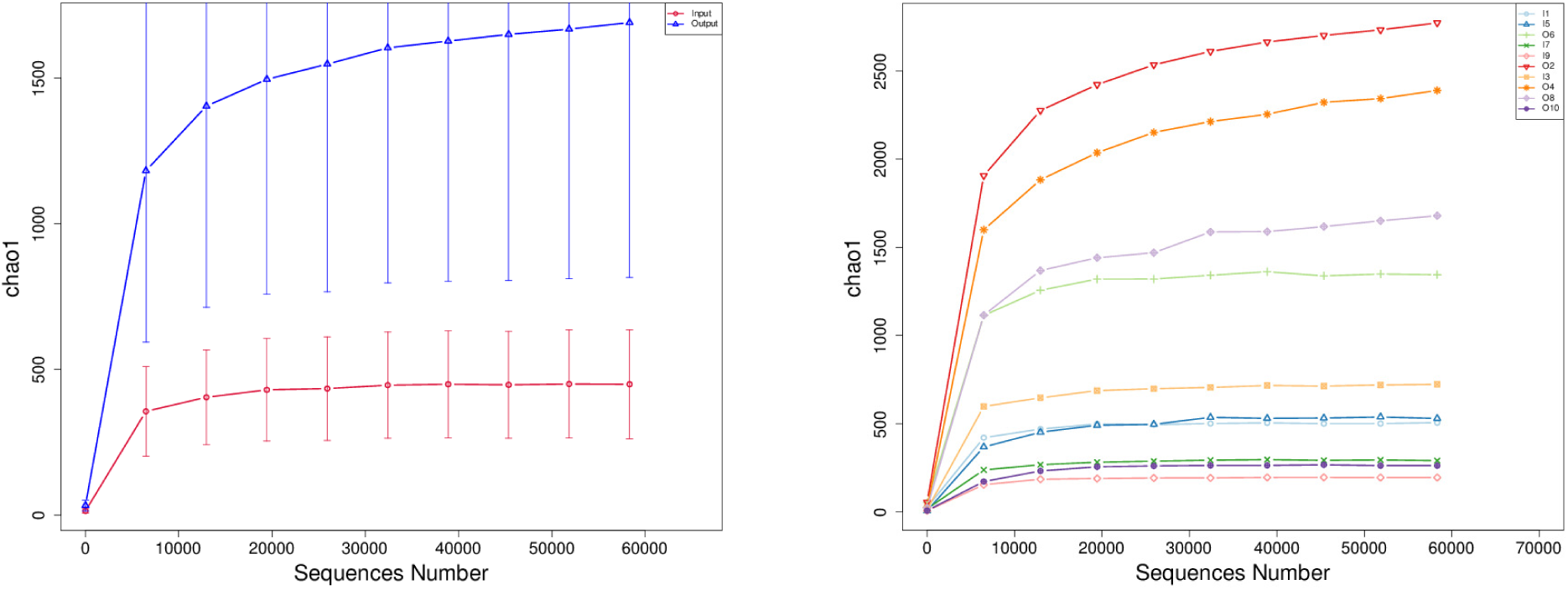
Chao1 richness rarefaction curves comparing alpha diversity across wastewater samples before (Input) and after (Output) treatment. (Right) Individual sample curves show that several Output samples, notably O2 and O4, achieve substantially higher species richness—exceeding 2,000 detected features—while Input samples plateau below 550. Error bars represent confidence intervals at each sequencing depth. (Left) Group-wise rarefaction curves for all samples highlight consistently greater richness in Output groups versus Input groups, with variation among effluents; most curves plateau, indicating sufficient sequencing effort. Together, these results demonstrate that wastewater treatment generally enhances microbial richness, though effects differ between effluent samples.

### Beta Diversity Analyses Reveal Consistent Community Restructuring

Multivariate analyses confirmed dramatic compositional turnover between influent and effluent microbiomes. Detrended correspondence analysis (DCA) and non-metric multidimensional scaling (NMDS), performed on phylogenetic (weighted/unweighted UniFrac) and compositional (Bray–Curtis) distances (Fig. 2), Principal Coordinates Analysis (PCoA) (Fig. A3), revealed robust and consistent separation between influent and effluent groups, with minimal within-group overlap. Effluent microbiomes clustered distinctly from influents along the primary DCA/NMDS axes, indicating strong treatment-driven shifts in overall community structure.

**Fig. 2.**
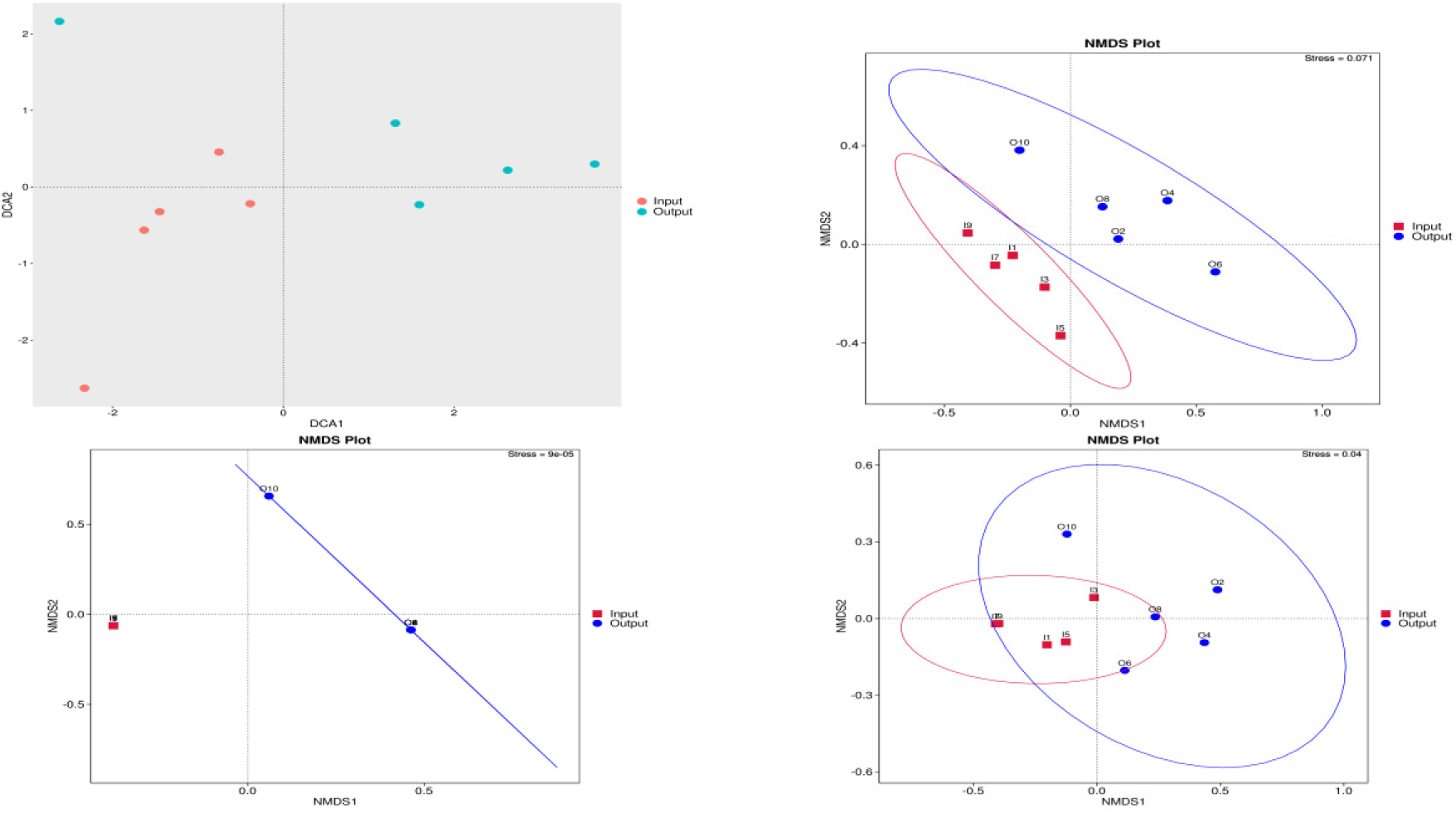
Detrended Correspondence Analysis (DCA) ordination plots depicting beta diversity of microbial communities before (Input) and after (Output) wastewater treatment demonstrate pronounced shifts in overall microbial community structure following wastewater treatment (Top-Left). Non-metric multidimensional scaling (NMDS) ordination plots illustrating beta diversity among influent (Input, red squares) and effluent (Output, blue circles) wastewater samples based on three distance metrics: weighted UniFrac (Down-left), unweighted UniFrac (Down-Right), and Bray–Curtis dissimilarity (Top-right). Each plot displays sample clustering along the primary NMDS axes, with 95% confidence ellipses outlining group dispersion. All distance metrics reveal clear separation between Input and Output communities, indicating that wastewater treatment induces pronounced and consistent shifts in both phylogenetic and compositional structure. Stress values indicate a robust fit for each ordination.

Statistical comparisons reinforced these findings: both weighted and unweighted UniFrac metrics indicated significantly greater within-group beta diversity among outputs (*t*-test weighted *p* = 0.0097, unweighted *p* = 0.0128; Wilcoxon *p* < 0.02 for both) (Fig. A4). These consistent results across multiple metrics highlight that treatment not only alters lineage abundance and dominance but also the fundamental membership of the microbial community.

Hierarchical clustering analysis (Fig. A5) with heatmaps of Bray–Curtis distances further visualized the emergence of two primary sample clusters strictly segregated by treatment stage. Pairwise distance heatmaps confirmed that between-group dissimilarities far exceeded within-group variations, a pattern also supported by 95% confidence ellipses on ordination plots (Fig. A6).

### Taxonomic Restructuring: From Dominance to Diversity

Taxonomic profiling at both the phylum and genus level revealed a transformation from a dominance-structured community in influents to a dramatically more even and taxonomically rich effluent microbiome (Fig. 3). Influent samples were almost universally dominated by Proteobacteria—primarily Pseudomonas (Gammaproteobacteria, Pseudomonadaceae)—with low overall richness and evenness. Following the treatment, the relative abundance of Proteobacteria declined markedly, while groups such as Actinobacteriota, Firmicutes, Bacteroidota, and Patescibacteria became prevalent. This taxonomic expansion in effluent samples was reflected at finer ranks by emergent or newly dominant genera, including Paenibacillus, Acinetobacter, Shewanella, Mycobacterium, Aeromonas, and Comamonas.

**Fig. 3.**
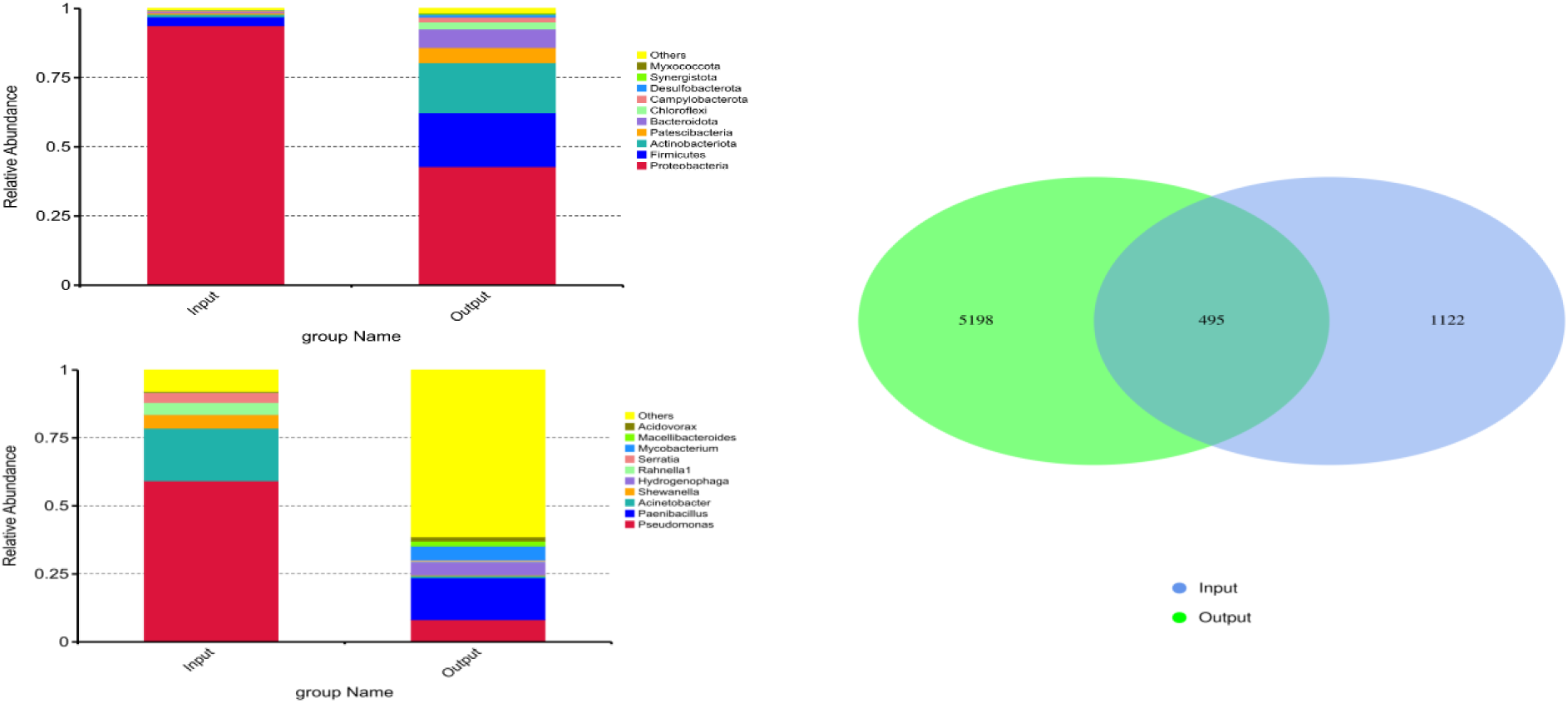
Stacked barplots comparing the relative abundances of dominant microbial taxa at the phylum level (Top-left) and genus level in influent (Input) and effluent (Output) wastewater samples (Down-Left). A Venn diagram illustrates the overlap of microbial taxa in influent (Input, blue) and effluent (Output, green) wastewater samples (Right).

Heatmap visualization of the 35 most abundant genera emphasized these community compositional changes (Fig. 4). Hierarchical clustering of genera showed clear segregation between influent-dominated (Pseudomonas, Escherichia) and effluent-enriched (Actinobacterium, Bacillus, Shewanella, Aeromonas, etc.) lineages, with effluents exhibiting greater intra-group taxonomic diversity (Fig. A7).

**Fig. 4.**
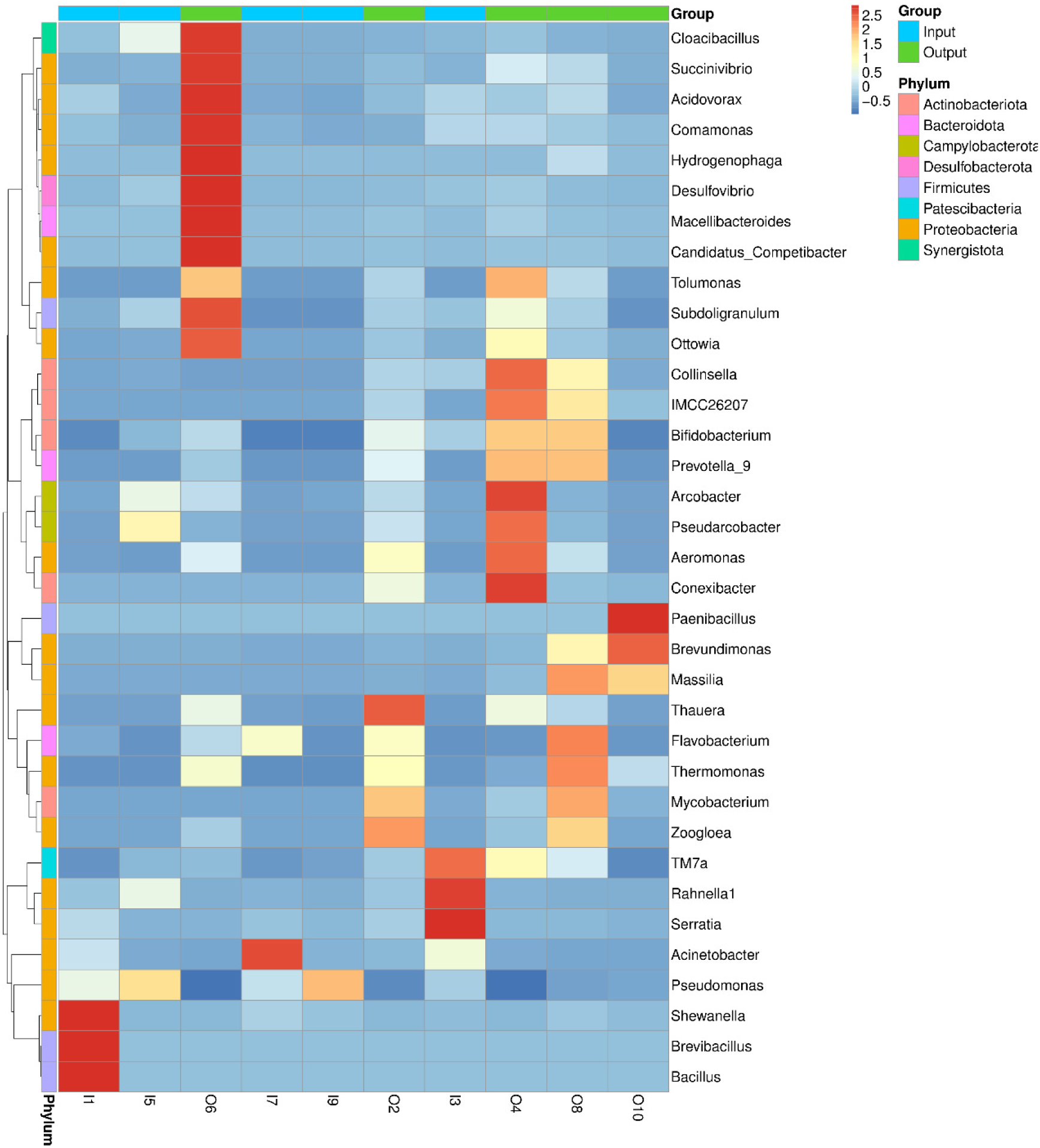
Heatmap displays the relative abundance of major bacterial genera across influent (Input) and effluent (Output) wastewater samples. Rows correspond to genera, color-coded by phylum (left bar), and columns to samples, grouped by treatment status (top bar). Color intensity indicates normalized abundance (blue: low, red: high). Clustering along both axes reveals group-specific and sample-specific shifts, with Output samples displaying increased diversity and enrichment of multiple genera, while certain genera are dominant in Input samples. These patterns highlight marked taxonomic restructuring of microbial communities following wastewater treatment.

A Venn diagram analysis revealed 495 ASVs shared by both groups, but 5,198 were exclusive to outputs and 1,122 to inputs. This indicates that treatment eliminates many influent-specific lineages and enables the emergence of novel effluent-associated taxa (Fig. 3).

### Persistence of Potential Pathogens in Treated Effluent

Despite the pronounced community restructuring and reduction in Proteobacteria dominance, clinically relevant lineages persisted in several treated samples. ASVs affiliated to Gammaproteobacteria (e.g., Escherichia, Pseudomonas, Legionella, Klebsiella, Campylobacter) remained common in some effluents; notably, Campylobacter reached relative abundances exceeding 3% in certain outputs, and multiple effluent samples (O2, O4, O8, O10) harbored non-trivial loads of taxa of known clinical concern.

This persistence underscores the public health risk posed by the partial survival or potential enrichment of opportunistic and enteric pathogens in treated wastewater, especially where water reuse or environmental release are common practices.

### Functional Potential: Adaptation, Redundancy, and Resilience

Predictive functional profiling revealed that both influent and effluent communities encoded a broad, highly redundant suite of metabolic and adaptive functions. Nevertheless, effluent communities displayed greater functional evenness, enrichment in nutrient cycling, xenobiotic degradation, and environmental stress adaptation functions, and increased beta diversity of predicted pathways. Key differentially abundant functions in effluents related to complex carbohydrate degradation, stress response, and environmental sensing, consistent with a microbiome adapted for oligotrophic, post-treatment conditions (Fig. 5).

**Fig. 5.**
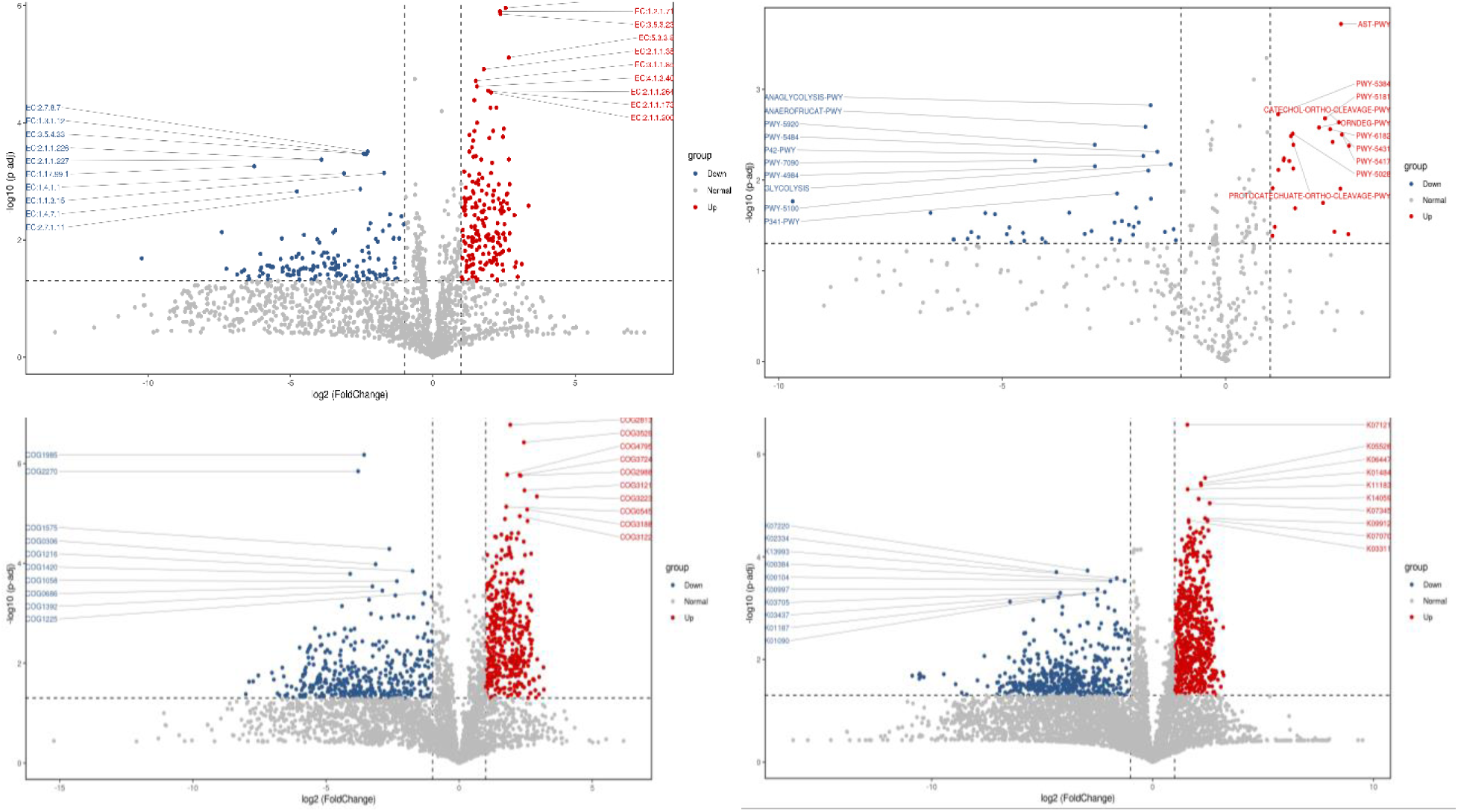
Grouped volcano plots illustrating the differential abundance of predicted microbial functions— (top left) EC numbers, (top right) pathways, (bottom left) COGs, and (bottom right) KEGG Orthologs (KOs)—between two wastewater groups. A volcano plot is a type of scatter plot that visually presents results of statistical comparisons, typically showing both significance and magnitude of change. In each panel, the horizontal axis displays the fold difference (log2 fold change) in abundance of each differential feature between groups, and the vertical axis represents the statistical significance of the between-group test (–log10 p-value). Each point corresponds to a specific functional feature: “Up” marks features with significantly higher abundance in the first group than in the second, while “Down” marks features with higher abundance in the second group. Red points highlight functions significantly enriched in the first group, blue points those enriched in the second, and gray points are not statistically different. Labeled outliers show the most strongly differentiated functions. This visualization reveals up- and down-regulated features at multiple functional levels, illustrating the extent and direction of functional shifts between groups.

## Discussion

WWTPs represent critical ecological filters, fundamentally transforming microbial communities while simultaneously tackling public health risks and environmental pollution. In the context of rapid urbanization and water scarcity, especially in arid regions like Egypt, understanding microbiome dynamics through comprehensive approaches—such as 16S rRNA deep sequencing and physicochemical analyses—is essential for sustainable water management and safe reuse strategies.

A foundational insight from this study is the consistent expansion of microbial richness and evenness in treated effluent relative to raw influent (39, 40). The ecological restructuring of wastewater microbiomes—from dominance-structured influents to high-richness, even effluents—is depicted in Fig. 6, visually summarizing the key findings of this study. Quantitative alpha diversity metrics—including Chao1, Shannon, and Simpson indices—reveal a nearly fourfold increase in richness post-treatment, with corresponding improvements in evenness and a reduction in dominance indices. Effluent rarefaction curves plateau at substantially higher ASV counts than any influent sample, signalling high sequencing depth and robust community complexity (Fig. 1).

**Fig. 6.**
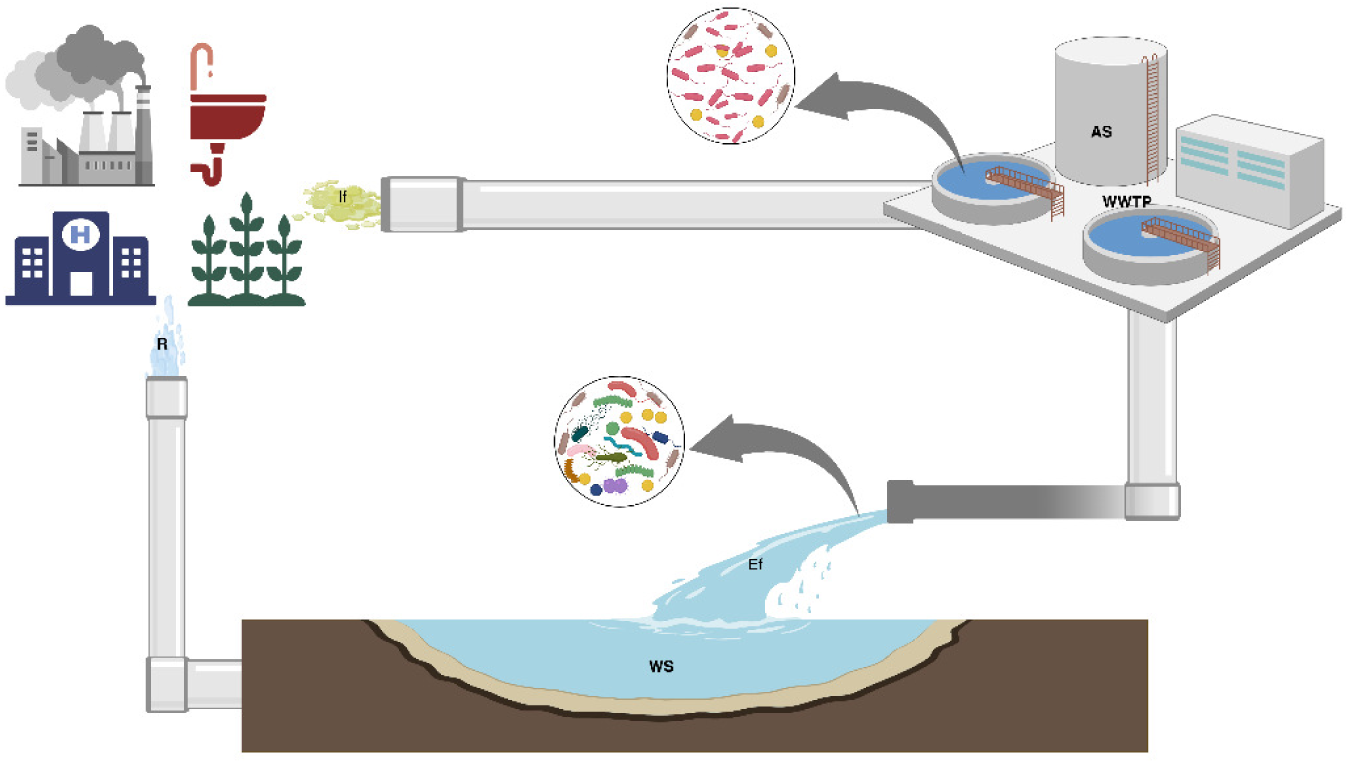
Schematic overview of microbial community restructuring during municipal wastewater treatment. Wastewater influent (If) from diverse urban and agricultural sources enters the wastewater treatment plant (WWTP) with activated sludge (AS) processing. The influent microbial community is dominance-structured, while the effluent (Ef) community released into the water stream (WS) displays increased richness and evenness with reduced dominance. Created in BioRender. Gamal, A. (2025) https://BioRender.com/jzw93x6.

This marked rise in diversity challenges the traditional view of WWTPs as mere kill steps for microorganisms. Instead, treatment acts as a selective ecological bottleneck, suppressing fast-growing, copiotrophic, and host-associated lineages (primarily Proteobacteria such as *Pseudomonas*) and fostering slow-growing, resilient, and functionally redundant taxa better adapted to nutrient-limited and dynamic post-treatment environments. This pattern mirrors broader ecological principles in disturbed natural and engineered ecosystems, wherein diversity and evenness often predict functional stability and ecological resilience.

Multivariate beta diversity analyses confirm a significant community restructuring. Methods such as DCA, NMDS, and PCoA consistently separate influent and effluent microbiomes across Bray-Curtis and UniFrac distance metrics (Fig. 2). Statistical frameworks (PERMANOVA, ANOSIM) attribute over 23% of community composition variance, highlighting the extent and reproducibility of these ecological shifts.

This restructuring, universal across all five WWTPs studied, is modulated by plant-specific influences including influent chemistry (e.g., elevated salinity in Plant 5) (Atalla *et. al.*, under review), operational modes, and catchment characteristics. Outliers in ordination space reflect the imprint of biogeographic and process adaptation factors, reinforcing the critical need to consider local context in interpreting microbiome outcomes.

Venn diagram analyses further underscore the non-attritional nature of WWTP microbial filtering, revealing thousands of unique effluent ASVs and novel taxa that emerge or proliferate post-treatment. This signals an ecological succession more akin to adaptation and niche expansion than simple reduction (Fig. 3).

Detailed taxonomic profiling demonstrates clear succession: influents are dominated by host-associated and fast-exploiting lineages such as *Pseudomonas*, *Escherichia*, and *Klebsiella*. Post-treatment, these are supplanted by stress-tolerant and metabolically versatile assemblages, including Actinobacteriota, Firmicutes, Bacteroidota, and Patescibacteria. At finer taxonomic tiers, genera like *Paenibacillus*, *Acinetobacter*, *Shewanella*, *Mycobacterium*, *Aeromonas*, and *Comamonas* emerge, each contributing specialized functions, biofilm formation, organic degradation, metal reduction, nutrient cycling, and stress tolerance (11, 41, 42).

Heatmap and clustering analyses further portray the occupation of novel ecological niches, a hallmark of successful engineering interventions and ecosystem resilience (Fig. 4).

Predictive functional profiling using COG, EC, KEGG ortholog, PFAM, and TIGRFAM annotations reveals broad metabolic capacity and pronounced redundancy in both influent and effluent communities. Effluents, however, display greater evenness in the distribution of metabolic pathways and increased capacity for nutrient cycling, xenobiotic degradation, and stress adaptation, marking an adaptive shift toward resilience in oligotrophic and chemically challenging conditions (Fig. 5).

Importantly, while no truly novel functions dominate the effluent, the greater evenness and redundancy suggest a distributed metabolic network, reinforcing the notion that functional resilience and reliability are enhanced by diversity and overlap in core capabilities. The identification of numerous “other” functions also highlights limitations in current annotation databases and underscores the need for future metagenomic and transcriptomic studies to fill gaps in functional understanding (43, 44).

Despite substantial reductions in classic sewage-associated and potentially harmful genera, several clinically relevant pathogens persist in treated effluents, notably *Escherichia*, *Pseudomonas*, *Klebsiella*, *Legionella*, and *Campylobacter*. The latter occasionally reaches relative abundances exceeding 3% in outputs, a level of concern given common standards of water reuse and environmental release.

The persistence of these and other risk genera indicates that conventional treatment is not always effective. Factors such as influent load, clarifier regrowth, plant design variables, and the inherent resistance of certain taxa may compromise complete pathogen removal. Moreover, the continued detection of genera associated with antibiotic resistance traits (e.g., *Escherichia*, *Klebsiella*, *Acinetobacter*) raises unresolved concerns about resistance dissemination in water systems and the overall effectiveness of WWTPs as barriers to the spread of antimicrobial resistance, a core concern under “One Health” frameworks demanding integrated human, animal, and environmental health surveillance.

There is a strong correlation between microbial changes and major physicochemical transitions effected by treatment. Removal of labile organic matter (COD, BOD, TOC) coincides with the decline of copiotrophs. TN and TP removal are linked to the enrichment of specialist denitrifiers (Rhodocyclales, Comamonadaceae) and phosphate-cycling taxa (Burkholderiales, Actinobacteria), respectively.

Site-to-site variability (e.g., TP increases in Plant 3; reduced TN removal in Plant 4) underpins the critical role of process configuration, operational best practices, and context-dependent influent chemistries in determining ecological outcomes. These results advocate for tailored optimizations, adaptive management, and targeted interventions rather than blanket solutions.

A major area of agreement among recent reports, including the present work and studies by Azli *et al.* (2022), Makuwa *et al.* (2023a), Oluseyi Osunmakinde *et al.* (2019), Tyagi *et al.* (2025), and Wardi *et al.* (2023), is the predominance of Proteobacteria in raw sewage and their pronounced reduction following biological treatment (11, 23, 26, 39, 40). This phylum’s dominance in influent is a globally recognized signature of human and animal waste streams. In parallel, most comparative studies, including those by Azli *et al.* (2022) and Masrahi (2023), consistently report post-treatment enrichment of Actinobacteriota, Firmicutes, and Bacteroidota in effluents (39, 45). Such transitions suggest strong selective pressures favoring stress-tolerant, metabolically versatile taxa during engineered treatment, supporting a core ecological paradigm of succession from copiotrophic, dominance-structured consortia toward more balanced, resilient microbial communities. Further, several studies partially mirror the current findings regarding the persistence of Bacteroidetes and the enrichment of Actinobacteriota and Firmicutes post-treatment (11, 26, 40, 46–48). Such results reinforce the broad ecological trends of bacterial community restructuring as a hallmark of modern wastewater treatment.

Crucially, the observation of effluent-resident genera with public health importance, such as *Escherichia*, *Legionella*, and *Campylobacter*, is also echoed in studies from Makuwa *et al.* (2023a), Numberger *et al.* (2019), and Wardi *et al.* (2023) (11, 23, 29). This persistent detection of clinically relevant and opportunistic pathogens, even in treated waters, underscores the partial limitations of current conventional treatment for complete risk abatement and aligns with international concerns about the environmental dissemination of potentially pathogenic taxa. Despite these patterns of agreement, notable divergences exist among microbial profiles reported after treatment. For example, Oluseyi Osunmakinde *et al.* (2019) observed effluent communities still dominated by Proteobacteria, Actinobacteria, Firmicutes, and Chloroflexi, with minimal contributions from Bacteroidota or Patescibacteria, diverging from our results where increased evenness and taxonomic expansion were prominent. Similarly, studies in wastewater-irrigated soils (Masrahi, 2023) documented continual dominance of Firmicutes, Actinobacteria, Proteobacteria, and Bacteroidetes without major shifts in phylum proportions, contrasting with the pronounced gain in evenness and diversity seen here (40, 45). There are also conflicting findings regarding the relative abundance of Proteobacteria in effluent: Tyagi *et al.* (2025) reported an increase rather than a decrease in Proteobacteria after treatment, while Wardi *et al.* (2023) described persistent co-dominance of Bacteroidetes and Proteobacteria from influent through effluent, with only a limited rise in pathogen frequency in one plant(11, 26). These divergences highlight how WWTP configuration, catchment-specific influent composition, geographic and climatic factors, and methodological heterogeneity, such as differences in sequencing depth, region of the 16S rRNA gene targeted, or sample handling, can all modulate post-treatment microbial structure. In terms of pathogenic diversity, broader spectra of medically relevant genera have been reported elsewhere: Oluseyi Osunmakinde *et al.* (2019) found widespread dissemination of genera like *Roseomonas*, *Bacillus*, *Clostridium*, *Mycobacterium*, *Methylobacterium*, and *Aeromonas*, while Makuwa *et al.* (2023a) documented detection of additional pathogens such as *Pseudomonas, Staphylococcus, Streptococcus*, *Vibrio, Leptospira, Salmonella, Yersinia, and Shigella*. In contrast, this study primarily identified *Escherichia, Legionella,* and *Campylobacter* within the effluent pathogen profile, suggesting plant and region-specific variation in pathogen persistence and risk (23, 40). Interpretation and Broader Implications Taken together, these comparative insights illuminate a significant foundation of ecological agreement and meaningful variation across studies. The pre-treatment dominance of Proteobacteria is almost universal, but the degree of post-treatment diversification, the relative abundances of dominant phyla, and the richness of effluent-associated pathogens are contingent on nuanced local determinants, plant technology, operational practices, influent source diversity, and the specific regulatory and reuse context. The observed heterogeneity underscores the importance of not extrapolating single-plant, single-region findings to broader contexts without careful consideration of environmental, industrial, and methodological variables. It also highlights the vital need for standardized, longitudinal, and multi-sample strategies, ideally tracing individual wastewater streams through treatment and into receiving environments, to more accurately capture treatment-driven microbial transformations and their ecological and public health risks. Ultimately, while wastewater treatment reliably shapes bacterial community structure, the trajectory and outcome of this restructuring are not invariant. The interplay of local conditions, plant design, and broader geographic context determines the degree to which microbiomes are “reset” during treatment, the functional redundancy preserved, and the residual public health risk for downstream water reuse. This plurality challenges us to design surveillance and intervention strategies that are both evidence-based and context-sensitive, positioning microbial ecology as central to sustainable wastewater.

These findings highlight the need for ongoing, nuanced surveillance of WWTP microbiomes, not just to verify removal efficiency, but to track community dynamics, functional stability, and public health risk. Integrating routine microbiome and resistome profiling into plant operations and regulations is essential to anticipate and prevent the discharge of persistent pathogens and resistance genes. Optimizing processes through advanced treatment, source-separated streams, and responsive operational adjustments will further support safe water reuse.

Future work should expand to include time series, activated sludge, and biofilm samples to clarify the mechanisms of community change and resilience. Region-specific, scalable monitoring and better integration of chemical-microbiological risk assessments—aligned with One Health—are critical for advancing water security in Egypt and comparable regions.

Key limitations remain: the lack of activated sludge data, single-time-point sampling, and the functional limits of 16S rRNA predictions. Whole metagenome/transcriptome studies are needed to resolve unknowns in function and risk.

In conclusion, Egyptian WWTPs reshape their incoming microbiomes from dominance by a few, potentially hazardous, copiotrophic lineages into diverse, functionally robust, yet occasionally pathogen-harboring effluent communities. This ecological renewal supports contaminant degradation and ecosystem resilience, but persistent pathogens require ongoing vigilance, technological refinement, and integrated monitoring for safe water reuse. As pressures on water resources escalate, leveraging microbial ecology for safer, more sustainable, and health-conscious wastewater management will be indispensable for national and global water security.

## Methods

### Study Design, Sample Collection, and Ethics

A cross-sectional study was conducted to characterize bacterial communities in influent and effluent samples from five representative WWTPs across Egypt. The selected WWTPs included: Balaqs (Al Qalyobia) (Latitudes: N 30°09’36.1” Longitudes: E 31°17’57.2”), Al-Berka (Cairo) (Latitudes: N 30°11’02.9” Longitudes E 31°24’56.0”), Nahtai (Al Gharbia, Nile Delta) (Latitudes: N 30°42’20.1” Longitudes: E 31°11’48.2”), Zenine (Giza) (Latitudes: N 30°01’59.3” Longitudes: E 31°10’57.8”), and Al Tanqya Sharqya (Alexandria, Nile Delta) (Latitudes: N 31°12’05.7” Longitudes: E 29°57’42.4”). Treatment systems predominantly employed conventional activated sludge, except Nahtai which utilizes a hybrid Moving Bed Biofilm Reactor (MBBR) and activated sludge process. Key microbial functional groups such as heterotrophic bacteria (removing organic loads measured as BOD and COD), nitrifying bacteria including ammonia-oxidizing bacteria (Nitrosomonas, Nitrosospira) and nitrite-oxidizing bacteria (Nitrospira, Nitrobacter) were identified as integral to system bioprocesses (49).

Sampling was conducted between October and November 2024 under approved institutional protocols with informed consent. At each plant, ten grab samples, 1 L influent and 5L effluent, were collected using sterilized plastic samplers following decontamination steps. Samples were immediately stored at 4°C and transported to Al-Dayora Central Laboratory, Greater Cairo Sanitary Drainage Company, for downstream analyses.

### Physicochemical wastewater quality assessment

In situ measurements of Temperature, pH, Dissolved Oxygen (DO), and Turbidity were obtained. Bulk parameters BOD, COD, TOC, TN, TP, and inorganic constituents (e.g., Total Dissolved Solids (TDS), Electrical Conductivity (EC), Cyanide (CN)) were assessed according to established protocols. Trace metals (Pb, Ni, Cd, Cu, Fe, Zn, Hg, As) were quantified by standard analytical methods (Atalla *et. al.*, under review).

### DNA Extraction and Molecular Processing

Influent samples underwent settling in Imhoff cones (WHEATON^®^) with sediment and supernatant fractions processed separately; effluent samples were filtered directly using sterile 0.45 μm membranes (Sartorious^®^), employing vacuum filtration systems (VACUUBRAND^®^). Samples were preserved at −20°C until DNA extraction, which was performed using combined biomass from sediment and membrane fractions. Biomass disruption was achieved by bead beating and heat treatment to ensure cell lysis. Genetic material was purified and quantified (Nanodrop). Its integrity is confirmed by agarose gel electrophoresis.

### 16S rRNA Gene Amplicon Sequencing

The extracted DNA samples were shipped to NOVOGENE^©^, China, for amplification, library construction, and next-generation sequencing. The PCR reaction was carried out with 15 μL of Phusion High-Fidelity PCR Master Mix, 0.2 μM of forward and reverse primers, and about 10 ng of template DNA. The V3–V4 region of the bacterial 16S rRNA gene was amplified using specific PCP primers: 341F (5′-CCTAYGGGRBGCASCAG-3′), and 806R (5′-GGACTACNNGGGTATCTAAT-3′). The thermal cycling consisted of initial denaturation at 98℃ for 1 min, followed by 30 cycles of denaturation at 98℃ for 10 s, annealing at 50℃ for 30 s, and elongation at 72℃for 30 s and 72℃ for 5 min. The PCR products were purified using magnetic bead purification. Samples were mixed in equi-density ratios based on the concentration of PCR products. After thorough mixing, the PCR products were detected, and the target bands were recovered. Sequencing libraries were generated, and indexes were added. The library was checked with Qubit and real-time PCR for quantification and a bioanalyzer for size distribution detection. Quantified libraries were pooled and sequenced on Illumina platforms, according to effective library concentration and the required data amount. Paired-end reads were assigned to samples based on their unique barcode and truncated by cutting off the barcode and primer sequence.

### Bioinformatics, functional prediction, and statistical data analysis

Paired-end reads were merged using FLASH (V1.2.1 1, http://ccb.jhu.edu/software/FLASH/) (50). A high-speed and accurate analysis tool was designed to merge paired-end reads when at least some of the reads overlap the read generated from the opposite end of the same DNA fragment, and the splicing sequences were called raw tags. Quality filtering on the raw tags was performed using the fastp (Version 0.23.1) software to obtain high-quality Clean Tags (51). The tags were compared with the reference database (Silva 138.1, https://www.arb-silva.de/) to detect chimera sequences, and the effective tags were obtained by removing the chimera sequences with the vsearch package (V2.16.0, https://github.com/torognes/vsearch) (52). For the Effective Tags obtained previously, denoise was performed with DADA2 in the QIIME2 software to obtain initial ASVs (53). For the obtained ASVs, on the one hand, the representative sequence of each ASV was annotated to obtain the corresponding species information and the abundance distribution based on species (54).

To study the phylogenetic relationship of each ASV and the differences of the dominant species among different samples (groups), multiple sequence alignment was performed using QIIME2 software. The absolute abundance of ASVs was normalized using a standard of sequence number corresponding to the sample with the fewest sequences. Subsequent analyses of alpha diversity and beta diversity were performed based on the output normalized data. The top 10 taxa of each sample at each taxonomic rank (Phylum, Class, Order, Family, Genus, Species) were selected to plot the distribution histogram of relative abundance in Perl through the SVG function. The abundance of the top 35 taxa of each sample at each taxonomic rank was used to draw the heatmap, which visually displays different abundance and taxa clustering. This was achieved in R through the pheatmap() function. Venn and Flower diagrams visually display the common and unique information between different samples or groups. Venn and Flower diagrams were produced in R with the VennDiagram() function and in Perl with the SVG function, respectively. A phylogenetic tree, also called an evolutionary tree, can describe the evolutionary relationship between different species. One hundred genera with the highest abundance in the samples were selected, and the sequence alignment was performed to draw the phylogenetic tree in Perl with the SVG function.

To analyze the diversity, richness, and uniformity of the communities in the sample, alpha diversity was calculated from 7 indices in QIIME2, including Observed_OTUs, Chao1, Shannon, Simpson, Dominance, Good’s coverage, and Pielou_e. Three indices were selected to identify community richness: Observed_OTUs – the number of observed species; Chao (http://scikit-bio.org/docs/latest/generated/skbio.diversity.alpha.observed_otus.html); – the Chao1 estimator (http://scikitbio.org/docs/latest/generated/skbio.diversity.alpha.chao1.html); Dominance – the Dominance index (http://scikitbio.org/docs/latest/generated/skbio.diversity.alpha.dominance.html); Two indices were used to identify community diversity: Shannon – the Shannon index (http://scikitbio.org/docs/latest/generated/skbio.diversity.alpha.shannon.html); Simpson – the Simpson index (http://scikit-bio.org/docs/latest/generated/skbio.diversity.alpha.simpson.html); One index was used to calculate sequencing depth: Coverage – the Good’s coverage (http://scikitbio.org/docs/latest/generated/skbio.diversity.alpha.goods_coverage.html); One index was used to calculate species evenness: Pielou_e – Pielou’s evenness index (http://scikitbio.org/docs/latest/generated/skbio.diversity.alpha.pielou_e.html). To evaluate the richness of the microbial community and sample size. The species accumulation boxplot can be used to visualize, which was performed with the vegan package in R software.

To evaluate the complexity of the community composition and compare the differences between samples (groups), beta diversity was calculated based on weighted and unweighted Unifrac distances in QIIME2. Beta diversity analysis was used to evaluate differences among samples in species complexity. Beta diversity on both weighted and unweighted Unifrac was calculated by QIIME software. Then, a heatmap was created to display the Unifrac distance between samples, which was realized in Perl. A cluster tree is constructed on UPGMA, which is based on the weighted unifrac distance matrix. This is widely used in ecology for evolutionary classification. UPGMA diagram was drawn through the upgma.tre () function within QIIME. Cluster analysis was preceded by PCA, which was applied to reduce the dimension of the original variables using the ade4 package and ggplot2 package with R software (Version 4.0.3). PCoA was performed to get principal coordinates and visualize complex and multidimensional data. A distance matrix of weighted or unweighted Unifrac among samples was obtained before transformation to a new set of orthogonal axes, by which the maximum variation factor is demonstrated by the first principal coordinate, and the second maximum one by the second principal coordinate, and so on. PCoA analysis was displayed by the ade4 package and the ggplot2 package in R software (Version 4.0.3). NMDS was also implemented for data dimension reduction. Like the PCoA, NMDS also uses the distance matrix, but it emphasizes the numerical rank instead. The distance between sample points on the diagram can only reflect the rank information rather than the numerical differences. NMDS analysis was implemented through R software with the ade4 package and the ggplot2 package.

A series of statistical analyses, which include Anosim, Adonis, Multi-response permutation procedure (MRPP), Simper, *t*-test, MetagenomeSeq, and LEfSe, were performed to reveal the community structure differentiation. Anosim, Adonis, and MRPP analyses are non-parametric tests that analyze the difference between high-dimensional data groups. They can test whether the differences between groups are significantly greater than the differences within the group, which can determine whether the grouping is meaningful. All of them were performed with the vegan package and the ggplot2 package within R. Simper can reveal the contribution of each species to the differentiation between groups. The top 10 species were selected and presented on the graph. It was performed in R with the Vegan package and the ggplot2 package. MetagenomeSeq can showcase the species that display significant differences between groups. It was performed in R with metagenomeSeq package. LEfSe is widely used to discover biomarkers, and it can reveal metagenomic characteristics. To achieve this, an exclusive package named lefse was utilized. The annotation results of the amplicons can also be associated with the corresponding functional databases, and the PICRUSt2 (V2.3.0) software can be used to predict and analyze the function of the microbial community in the ecological sample.

## Data Availability

16srRNA V3 and V4 amplicon sequencing data generated for this study have been archived in the NCBI Sequence Read Archive (SRA) under project accession PRJNA1310270.

## Code availability

https://github.com/Shaymaamalek/Supplementray_data.git

## Acknowledgement

Greater Cairo Sanitary Drainage Company, Drinking Water and Wastewater Company in Gharbia, Drinking Water and Wastewater Company in Giza, Alexandria Sanitary Drainage Company, and Al-Dayora Central Lab - GCSDC, for their support in collecting samples and chemical analysis.

Perplexity platform for refinement of the English language of this manuscript https://www.perplexity.ai/.

## Author Contributions

A.G.A. collected the samples, metadata, performed the physicochemical analysis, prepared the samples for DNA extraction, PCR, QC, Funding, Writing the manuscript, and revision.

E.M.I. helped the first author in determining the chemical parameters.

Sh. A. Principal Coordinator of the project, supervising the collection of the samples and preparing the DNA for PCR and sequencing, Data analysis, Manuscript writing, and reviewing.

## Competing interest

The authors declare no competing interests.

## Funding

The authors received no specific funding for this work.

## Abbreviations

(mg/L): Milligrams per Liter
(%): Percent
(16S rRNA): 16S ribosomal Ribonucleic acid
(V3–V4): Variable regions 3 and 4
(Q20): Phred quality score 20
(Q30): Phred quality score 30
(nt): Nucleotide
(L): Liter
(°C): degrees Celsius
(Pb): Lead
(Ni): Nickel
(Cd): Cadmium
(Cu): Copper
(Fe): Iron
(Zn): Zinc
(Hg): Mercury
(As): Arsenic
(μm): micrometer
(DNA): Deoxyribonucleic acid
(μL): microliter
(PCR): Polymerase Chain Reaction
(μM): micromolar
(ng): nanogram
(F): Forward primer
(R): Reverse primer
(s): Seconds
(min): Minutes
(QIIME2): Quantitative Insights Into Microbial Ecology 2
(Unifrac): Unique Fraction Metric
(UPGMA): Unweighted Pair Group Method with Arithmetic

## Appendix

**Table A1:** Removal efficiency (%) for 17 wastewater quality parameters across five treatment plants. Values indicate the percentage reduction from influent to effluent for each parameter and plant, summarizing overall treatment performance for physicochemical, organic, aggregate, and heavy metal indicators (Atalla et al., under review)..

| Sample | plant1 | plant2 | plant3 | plant4 | plant5 |
| --- | --- | --- | --- | --- | --- |
| Turbidity_eff | 98.26 | 96.8 | 98.32 | 84.62 | 95.77 |
| BOD5_eff | 94.5 | 92.59 | 99.42 | 98.02 | 79.35 |
| COD_eff | 94.59 | 98.62 | 91.03 | 92.62 | 89.18 |
| TOC_eff | 73.98 | 68.42 | 77.85 | 74.45 | 81.88 |
| TDS_eff | 7.18 | 10.63 | 16.45 | 1.28 | 8.06 |
| EC_eff | 7.35 | 7.75 | 14.78 | 1.78 | 15.74 |
| CN_eff | 40 | 95.65 | 82.22 | -100 | 62.5 |
| TP_eff | 31.16 | 17.55 | -21.79 | 31.84 | 8.33 |
| TN_eff | 60.28 | 55.86 | 60.94 | 41.03 | 54.71 |
| Pb_eff | 95.51 | 95.04 | 95.89 | 94.68 | 95.56 |
| Ni_eff | 99.57 | 99.71 | 99.67 | 99.78 | 99.8 |
| Cd_eff | 100 | 100 | 100 | 0 | 100 |
| Cu_eff | 100 | 100 | 100 | 100 | 100 |
| Fe_eff | 100 | 100 | 100 | 100 | 100 |
| Zn_eff | 100 | 2.97 | 100 | 100 | 100 |
| Hg_eff | 91.89 | 88.52 | 89.43 | 89.76 | 88.28 |
| As_eff | 84.21 | 100 | 60 | 94.44 | 100 |

**Fig. A2.**
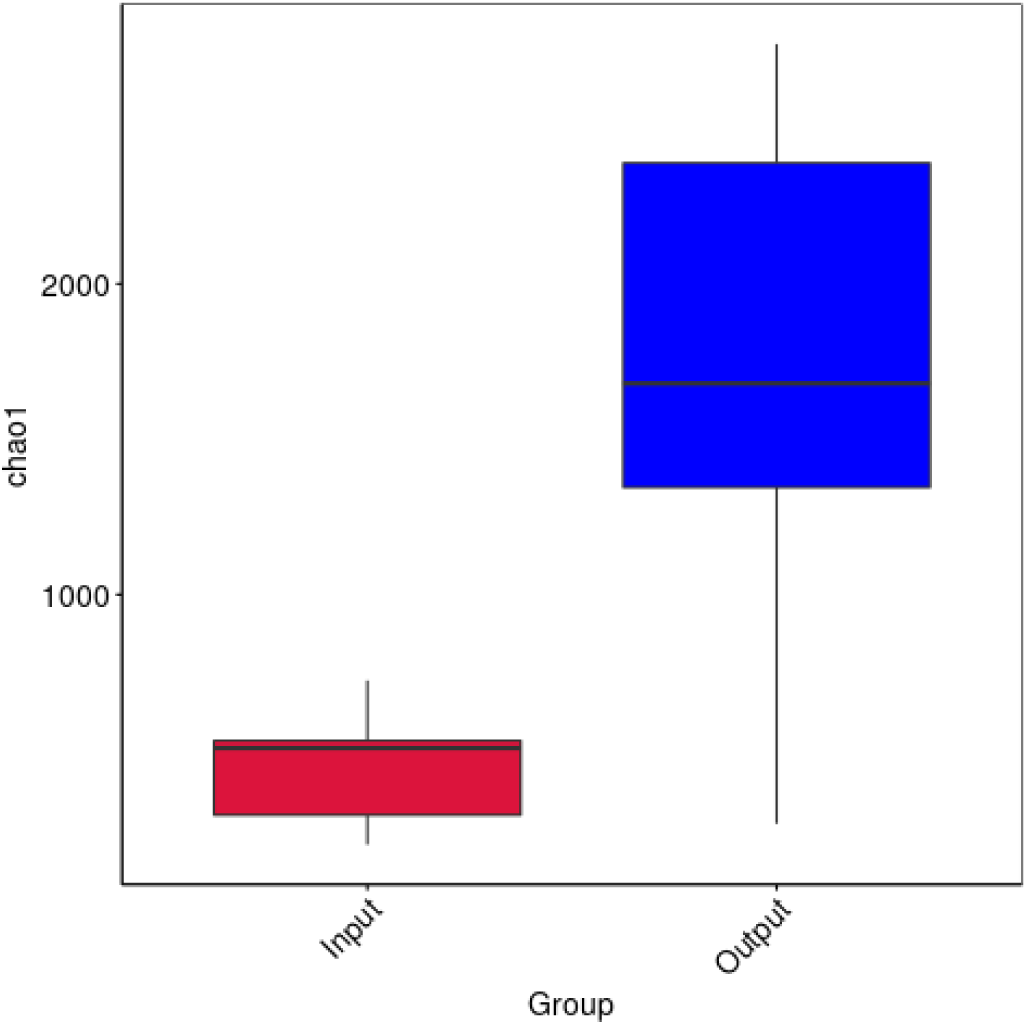
Chao1 Richness Boxplots with Statistical Test: Chao1 richness estimates for influent (Input, red) and effluent (Output, blue) wastewater samples. Effluent communities exhibit significantly greater richness (median ∼6,440 vs. ∼1,693 for influent; Wilcoxon p = 0.045), indicating substantial expansion of microbial diversity following treatment. Boxplots show median, interquartile range, and outliers.

Fig. A3 Principal Coordinates Analysis (PCoA) ordination plots showing beta diversity among influent (Input, red squares) and effluent (Output, blue circles) wastewater samples based on three distance metrics: weighted UniFrac (left), unweighted UniFrac (middle), and Bray–Curtis dissimilarity (right). Each panel illustrates sample distribution along the primary coordinate axes with 95% confidence ellipses for each group. Across all metrics, Input and Output communities form distinct, non-overlapping clusters, demonstrating that wastewater treatment leads to marked and consistent shifts in microbial community composition in both phylogenetic (presence/absence and abundance) and compositional structure. Axis labels indicate the proportion of variance explained by each principal coordinate.

Fig. A4 Comparison of beta diversity between influent (Input) and effluent (Output) wastewater samples using weighted and unweighted UniFrac distances. Boxplots depict pairwise distances for each group and metric: unweighted UniFrac (left, red and blue) and weighted UniFrac (right, red and blue). Output samples consistently exhibit significantly greater beta diversity than Inputs for both metrics, reflecting pronounced shifts in community structure following treatment. Statistical significance was confirmed by t-test (unweighted: p = 0.013; weighted: p = 0.0097) and Wilcoxon rank-sum test (unweighted: p = 0.019; weighted: p = 0.019). Together, these results indicate that wastewater treatment substantially alters both presence/absence-based and abundance-weighted phylogenetic community composition.

Fig. A5 Hierarchical clustering dendrograms and corresponding phylum-level stacked bar plots for influent (Input, red) and effluent (Output, blue) wastewater samples, using unweighted (right panels) and weighted (left panels) UniFrac distances. Dendrograms visualize phylogenetic composition, with weighted UniFrac reflecting abundance-weighted relationships and unweighted representing presence/absence patterns. In both approaches, Input and Output communities form distinct clusters, indicating marked shifts associated with treatment. Stacked bar plots display the relative abundance of major bacterial phyla for each sample, illustrating substantial changes in taxonomic composition and diversity between the two groups. Together, these panels demonstrate pronounced restructuring of microbial communities post-treatment at both phylogenetic and taxonomic levels.

Fig. A6 Heatmaps depicting pairwise beta diversity among wastewater samples (right) and sample groups (left) before (Input) and after (Output) treatment. In each matrix, the upper triangle shows weighted UniFrac distances (abundance- and phylogeny-based), while the lower triangle displays unweighted UniFrac distances (presence/absence-based). Color intensity ranges from red (low dissimilarity) to yellow (high dissimilarity), as indicated by the color bar. Both weighted and unweighted metrics reveal greater beta diversity between Input and Output groups than within groups, with weighted UniFrac capturing additional compositional differences driven by taxa abundance shifts. Together, these heatmaps illustrate that wastewater treatment induces pronounced and consistent restructuring of microbial communities at both the group and individual sample levels.

Fig. A7 Differentially abundant microbial taxa distinguishing influent (Input, red) and effluent (Output, green) wastewater communities, as identified by LEfSe analysis. (Left) Taxonomic cladogram displaying the phylogenetic relationships of taxa significantly enriched in each group. Node colors correspond to group enrichment: green for Output-associated lineages (notably within Actinobacteriota and Chloroflexi) and red for Input-associated taxa (primarily Gammaproteobacteria and related families). (Right) Barplot of LDA scores for taxa discriminatory between groups, highlighting the pronounced depletion of Pseudomonadaceae, Moraxellaceae, and other Gammaproteobacteria in effluent, while Output samples are enriched for diverse Actinobacteriota, Chloroflexi, and related clades. Together, these results demonstrate a marked shift in both the composition and phylogenetic structure of wastewater microbiota following treatment.

## Notes

### Competing Interest Statement

The authors have declared no competing interest.

## References

1. Unesco. 2023. Wastewater: the new black gold

2. UN Water. 2021. Progress on Wastewater Treatment – 2021 Update.

3. Varma VKC, Rathinam R, Suresh V, Naveen S, Satishkumar P, Abdulrahman IS, Salman HM, Singh P, Kumar JA. 2024. Urban waste water management paradigm evolution: Decentralization, resource recovery, and water reclamation and reuse. Environmental Quality Management 33:523–540.

4. Alagha O, Allazem A, Bukhari AA, Anil I, Mu’azu ND. 2020. Suitability of SBR for Wastewater Treatment and Reuse: Pilot-Scale Reactor Operated in Different Anoxic Conditions. Int J Environ Res Public Health 17:1617.

5. Villarín MC, Merel S. 2020. Paradigm shifts and current challenges in wastewater management. J Hazard Mater 390:122139.

6. Tortajada C. 2020. Contributions of recycled wastewater to clean water and sanitation Sustainable Development Goals. NPJ Clean Water 3:22.

7. WHO. 2006. Guidelines for the safe use of wastewater, excreta and greywater in agriculture and aquaculture. World Health Organization.

8. Shoushtarian F, Negahban-Azar M. 2020. World wide regulations and guidelines for agriculturalwater reuse: A critical review. Water (Switzerland). MDPI AG 10.3390/W12040971.

9. United Nations General Assembly (UNGA). 2015. 70/1. Transforming our world: the 2030 Agenda for Sustainable Development Transforming our world: the 2030 Agenda for Sustainable Development Preamble.

10. UN Water. 2017. Wastewater the untapped resources.

11. Wardi M, Slimani N, Alla AA, Belmouden A. 2023. First study of the effect of wastewater treatment on microbial biodiversity at three wastewater treatment plants in Agadir, Morocco, using 16S rRNA sequencing. Environmental Pollution 337:122528.

12. Duque AF, Campo R, Rio AV Del, Amorim CL. 2021. Wastewater valorization: Practice around the world at pilot-and full-scale. Int J Environ Res Public Health. MDPI 10.3390/ijerph18189466.

13. Naidoo S, Olaniran AO. 2013. Treated wastewater effluent as a source of microbial pollution of surface water resources. Int J Environ Res Public Health. MDPI 10.3390/ijerph110100249.

14. Topaz T, Egozi R, Suari Y, Ben-Ari J, Sade T, Chefetz B, Yahel G. 2020. Environmental risk dynamics of pesticides toxicity in a Mediterranean micro-estuary. Environmental Pollution 265:114941.

15. Ramírez-Morales D, Pérez-Villanueva ME, Chin-Pampillo JS, Aguilar-Mora P, Arias-Mora V, Masís-Mora M. 2021. Pesticide occurrence and water quality assessment from an agriculturally influenced Latin-American tropical region. Chemosphere 262:127851.

16. Rajmohan KS, Chandrasekaran R, Varjani S. 2020. A Review on Occurrence of Pesticides in Environment and Current Technologies for Their Remediation and Management. Indian J Microbiol 60:125–138.

17. Khan A, Ali J, Jamil SUU, Zahra N, Tayaba TB, Iqbal MJ, Waseem H. 2022. Removal of micropollutants, p. 443–461. In Environmental Micropollutants. Elsevier.

18. Wu K, Atasoy M, Zweers H, Rijnaarts H, Langenhoff A, Fernandes T V. 2023. Impact of wastewater characteristics on the removal of organic micropollutants by Chlorella sorokiniana. J Hazard Mater 453:131451.

19. Luo Y, Guo W, Ngo HH, Nghiem LD, Hai FI, Zhang J, Liang S, Wang XC. 2014. A review on the occurrence of micropollutants in the aquatic environment and their fate and removal during wastewater treatment. Science of The Total Environment 473–474:619–641.

20. Lee B, Lee S-H, Lee J, Kim N-K, Kim I, Park S, Kim S, Ham S-Y, Park H-D, Im D, Kim H-S. 2025. Influence of wastewater type on the distribution of microbial community compositions including pathogenic bacteria within wastewater treatment processes. Sustainable Environment Research 35:17.

21. Sasi R, Suchithra T V. 2023. Wastewater microbial diversity versus molecular analysis at a glance: a mini-review. Brazilian Journal of Microbiology 54:3033– 3039.

22. Kalmakhanova MS, Diaz de Tuesta JL, Malakar A, Gomes HT, Snow DD. 2023. Wastewater Treatment in Central Asia: Treatment Alternatives for Safe Water Reuse. Sustainability 15:14949.

23. Makuwa S, Green E, Fosso-Kankeu E, Moroaswi V, Tlou M. 2023. A Snapshot of the Influent and Effluent Bacterial Populations in a Wastewater Treatment Plant in the North-West Province, South Africa. Appl Microbiol 3:764–773.

24. Iloms EC. 2024. Microbial Diversity in Wastewater Sources and Biological Activated Sludge System. Int J Agric Biol 33:330310–330310.

25. Yasir M. 2020. Analysis of Microbial Communities and Pathogen Detection in Domestic Sewage Using Metagenomic Sequencing. Diversity (Basel) 13:6.

26. Tyagi I, Tyagi K, Ahamad F, Bhutiani R, Kumar V. 2025. Assessment of Bacterial Community Structure, Associated Functional Role, and Water Health in Full-Scale Municipal Wastewater Treatment Plants. Toxics 13.

27. Selvarajan R, Sibanda T, Pandian J, Mearns K. 2021. Taxonomic and Functional Distribution of Bacterial Communities in Domestic and Hospital Wastewater System: Implications for Public and Environmental Health. Antibiotics 10:1059.

28. Xie N, Zhong L, Ouyang L, Xu W, Zeng Q, Wang K, Zaynab M, Chen H, Xu F, Li S. 2021. Community composition and function of bacteria in activated sludge of municipal wastewater treatment plants. Water (Switzerland) 13.

29. Numberger D, Ganzert L, Zoccarato L, Mühldorfer K, Sauer S, Grossart H-P, Greenwood AD. 2019. Characterization of bacterial communities in wastewater with enhanced taxonomic resolution by full-length 16S rRNA sequencing. Sci Rep 9:9673.

30. Singh S, Ahmed AI, Almansoori S, Alameri S, Adlan A, Odivilas G, Chattaway MA, Salem S Bin, Brudecki G, Elamin W. 2024. A narrative review of wastewater surveillance: pathogens of concern, applications, detection methods, and challenges. Front Public Health 12.

31. Riesenberger B, Rodriguez M, Marques L, Cervantes R, Gomes B, Dias M, Pena P, Ribeiro E, Viegas C. 2024. Filling the Knowledge Gap Regarding Microbial Occupational Exposure Assessment in Waste Water Treatment Plants: A Scoping Review. Microorganisms. Multidisciplinary Digital Publishing Institute (MDPI) 10.3390/microorganisms12061144.

32. Central Agency for Public Mobilization and Statistics (CAPMAS). 2024. Annual Bulletin for Water Purification and Sanitation Statistics Year. Cairo, Egypt.

33. Shaheen MNF, Elmahdy EM, Rizk NM, Abdo SM, Hussein NA, Elshershaby A, Shahein YE, Fawzy ME, El-Liethy MA, Marouf MA, Abdel-Gawad FKh, Hu A, Gad M. 2024. Evaluation of physical, chemical, and microbiological characteristics of waste stabilization ponds, Giza, Egypt. Environ Sci Eur 36:170.

34. El-Feky AM, Saber M, Abd-El-Kader MM, Kantoush SA, Sumi T, Alfaisal F, Abdelhaleem A. 2024. Comprehensive environmental impact assessment and irrigation wastewater suitability of the Arab El-Madabegh wastewater treatment plant, ASSIUT CITY, EGYPT. PLoS One 19.

35. Osman HEM, Abdel-Hamed EMW, Al-Juhani WSM, Al-Maroai YAO, El-Morsy MHE-M. 2021. Bioaccumulation and human health risk assessment of heavy metals in food crops irrigated with freshwater and treated wastewater: a case study in Southern Cairo, Egypt. Environmental Science and Pollution Research 28:50217–50229.

36. Ayoub M, El-Morsy A. 2021. Applying the Wastewater Quality Index for Assessing the Effluent Quality of Recently Upgraded Meet Abo El-koum Wastewater Treatment Plant. Journal of Ecological Engineering 22:128–133.

37. Mohmed AH, Hameed A, El-Aassar M, Shawky HA, Mehany MAS, Khalil MMH. 2025. Assessment of Wastewater Treatment Plants (WWTPs) Performance in El-Sharkia, Egypt.

38. Zaky T, Abdelgawad H, Fatah M, Gamal A. 2022. Performance Evaluation of Wastewater Treatment Plants in El-Fayoum. International Journal of Environmental Studies and Researches 1:46–54.

39. Azli B, Razak MN, Omar AR, Mohd Zain NA, Abdul Razak F, Nurulfiza I. 2022. Metagenomics Insights Into the Microbial Diversity and Microbiome Network Analysis on the Heterogeneity of Influent to Effluent Water. Front Microbiol 13.

40. Oluseyi Osunmakinde C, Selvarajan R, Mamba BB, Msagati TAM. 2019. Profiling Bacterial Diversity and Potential Pathogens in Wastewater Treatment Plants Using High-Throughput Sequencing Analysis. Microorganisms 7:506.

41. Acharya K, Halla FF, Massawa SM, Mgana SM, Komar T, Davenport RJ, Werner D. 2020. Chlorination effects on DNA based characterization of water microbiomes and implications for the interpretation of data from disinfected systems. J Environ Manage 276:111319.

42. Tiwari A, Hokajärvi A-M, Domingo JS, Elk M, Jayaprakash B, Ryu H, Siponen S, Vepsäläinen A, Kauppinen A, Puurunen O, Artimo A, Perkola N, Huttula T, Miettinen IT, Pitkänen T. 2021. Bacterial diversity and predicted enzymatic function in a multipurpose surface water system – from wastewater effluent discharges to drinking water production. Environ Microbiome 16:11.

43. Douglas GM, Maffei VJ, Zaneveld JR, Yurgel SN, Brown JR, Taylor CM, Huttenhower C, Langille MGI. 2020. PICRUSt2 for prediction of metagenome functions. Nat Biotechnol 38:685–688.

44. Sun S, Jones RB, Fodor AA. 2020. Inference-based accuracy of metagenome prediction tools varies across sample types and functional categories. Microbiome 8:46.

45. Masrahi AS. 2023. Effect of long-term influx of tertiary treated wastewater on native bacterial communities in a dry valley topsoil: 16S rRNA gene-based metagenomic analysis of composition and functional profile. PeerJ 11.

46. Cai L, Ju F, Zhang T. 2014. Tracking human sewage microbiome in a municipal wastewater treatment plant. Appl Microbiol Biotechnol 98:3317–3326.

47. García-Armisen T, İnceoğlu Ö, Ouattara NK, Anzil A, Verbanck MA, Brion N, Servais P. 2014. Seasonal Variations and Resilience of Bacterial Communities in a Sewage Polluted Urban River. PLoS One 9:e92579.

48. Silva-Bedoya LM, Sánchez-Pinzón MS, Cadavid-Restrepo GE, Moreno-Herrera CX. 2016. Bacterial community analysis of an industrial wastewater treatment plant in Colombia with screening for lipid-degrading microorganisms. Microbiol Res 192:313–325.

49. Metcalf & Eddy Inc. 2013. Wastewater Engineering: Treatment and Resource Recovery, 5th ed. McGraw-Hill Education.

50. Magoč T, Salzberg SL. 2011. FLASH: fast length adjustment of short reads to improve genome assemblies. Bioinformatics 27:2957–2963.

51. Bokulich NA, Subramanian S, Faith JJ, Gevers D, Gordon JI, Knight R, Mills DA, Caporaso JG. 2013. Quality-filtering vastly improves diversity estimates from Illumina amplicon sequencing. Nat Methods 10:57–59.

52. Edgar RC, Haas BJ, Clemente JC, Quince C, Knight R. 2011. UCHIME improves sensitivity and speed of chimera detection. Bioinformatics 27:2194–2200.

53. Li M, Shao D, Zhou J, Gu J, Qin J, Chen W, Wei W. 2020. Signatures within esophageal microbiota with progression of esophageal squamous cell carcinoma. Chinese Journal of Cancer Research 32:755–767.

54. Callahan BJ, McMurdie PJ, Holmes SP. 2017. Exact sequence variants should replace operational taxonomic units in marker-gene data analysis. ISME J 11:2639–2643.

